# Bilayer-forming lipids enhance archaeal monolayer membrane stability

**DOI:** 10.1101/2025.02.20.639260

**Authors:** M. Saracco, P. Schaeffer, M. Tourte, S.-V. Albers, Y. Louis, J. Peters, B. Demé, S. Fontanay, P. Oger

**Affiliations:** INSA Lyon, Universite Claude Bernard Lyon 1, CNRS UMR5240, Villeurbanne, France; University of Strasbourg, CNRS, UMR 7177, F-67000 Strasbourg, France; University of Freiburg, Institute of Biology, Molecular Biology of Archaea, Freiburg, Germany; Institut Laue Langevin, Large Scale Structures group, Grenoble F-38042, France; Université Grenoble Alpes, CNRS, LiPhy, 38400 Grenoble, France; Institut Universitaire de France, 75231 Paris, France

## Abstract

Archaeal membranes exhibit remarkable stability under extreme environmental conditions, a feature attributed to their unique lipid composition. While it is widely accepted that tetraether lipids confer structural integrity by forming monolayers, the role of bilayer-forming diether lipids in membrane stability remains unclear. Here, we demonstrate that the incorporation of diethers into archaeal-like lipid assemblies enhances membrane organization and adaptability under thermal stress. Using neutron diffraction, we show that membranes composed of mixed diethers and tetraethers exhibit greater structural order and stability compared to pure lipid systems. Contrary to expectations, monolayer-forming tetraethers alone display increased variability in lamellar spacing under fluctuating temperature and humidity, whereas mixed lipid membranes maintain a consistent architecture. Furthermore, neutron scattering length density profiles reveal an unexpected density feature at the bilayer midplane, challenging conventional models of archaeal monolayer organization. These findings suggest that molecular diversity of lipid molecules, rather than tetraether dominance, plays a critical role in membrane auto assembly, stability and adaptability. Our results provide new insights into archaeal membrane adaptation strategies, with implications for the development of bioinspired, robust synthetic membranes for industrial and biomedical applications.

## Introduction

All living cells synthesize one or more plasma membranes, which isolate intracellular space from the external environment. Acting as a diffusion barrier for ions and small solutes, the membrane is essential not only for maintaining cellular integrity but also as a support for numerous cellular activities. This dynamic structure controls cell shape and growth, mediates vesicle release and integration, and facilitates intercellular communication (Konings et al., 2002; Shi et al., 2018).

Among the domains of life, Archaea are distinguished by the unique composition of their membrane lipids, setting them apart from Bacteria and Eukarya. Archaeal lipids are characterized by two polyisoprenoid alkyl chains linked to glycerol via ether bonds in a *sn*-2,3 configuration (Zhang, 1990). In contrast, bacterial and eukaryotic membrane lipids comprise two fatty acyl chains ester-linked in a *sn*-1,2 configuration (Nam et al., 2020).

These distinct features have been characterized as successful adaptations, enabling Archaea to thrive in extreme environments. The ether linkages in archaeal membranes are more chemically and thermally resistant than the ester linkages in bacterial and eukaryotic membranes. Furthermore, the ether bonds in the hydrophobic core promote tighter packing of polar headgroups, increasing water impermeability and reducing electrostatic interactions (Kruczek et al., 2017; Mathai et al., 2001). In addition, the archaeal isoprenoid hydrocarbon chains with branched methyl groups enhance lipid packing and membrane rigidity compared to the linear acyl chains generally found in bacterial and eukaryotic lipids. These adaptations result in increased stability and impermeability under extreme conditions (Komatsu and Chong, 1998; Polak et al., 2014).

Moreover, Archaea can synthesize monolayer-forming tetraether lipids (GDGTs), which consist of two C_40_ isoprenoid side chains ether-linked to two glycerol moieties (Boyd et al., 2013). The structural diversity of tetraether lipids includes glycerol mono-, di-, and trialkyl glycerol tetraethers (GMGT, GDGT, and GTGT, respectively) (Knappy et al., 2011), with unsaturated/cyclized, hydroxylated, or methylated isoprenoid chains (Nichols et al., 2004). Tetraether lipids may also contain up to eight cyclopentane rings and/or one cyclohexane ring (Zeng et al., 2019). These bipolar lipids exhibit unique behaviors, forming membranes that are more stable, rigid, and impermeable than those of Bacteria or Eukarya (Komatsu and Chong, 1998). By forming monolayer membranes, lipid packing is increased, aiding the regulation of solute and proton fluxes across the membrane (Cario et al., 2015; Matsuno et al., 2009). These extraordinary membrane lipid compositions establish Archaea as true champions of extreme environments, outperforming other organisms in nearly all environmental conditions (Rampelotto, 2010).

Archaea are particularly adapted to growth at high temperatures, with *Methanopyrus kandleri* holding the record for growing at 122 °C (Takai et al., 2008). The physicochemical properties of archaeal lipids, as described above, equip their membranes for stability in extremely hot habitats. At high temperatures, membranes typically experience increased fluidity and permeability. To compensate, bacteria and eukaryotes adjust the length, saturation, and branching of hydrophobic chains and the proportion of specific polar headgroups (Jebbar et al., 2015; Siliakus et al., 2017). To maintain physiological homeostasis, membrane integrity is continuously secured through homeoviscous adaptation, a fundamental mechanism in which membrane lipid composition is dynamically adjusted to maintain optimal fluidity and functionality (Hazel, 1995; Sinensky, 1974). In Archaea, homeoviscous adaptation involves distinct mechanisms. For example, cyclopentane rings are introduced into hyperthermophilic archaeal membranes to enhance rigidity, as observed in *Sulfolobus acidocaldarius* (T_max_ = 90 °C) and *Thermoplasma acidophilum* (T_max_ = 60 °C) (Uda et al., 2001; Zhou, 2020). Other, heat-adapted membranes show an increased proportion of macrocyclic tetraethers (GMGT), as well as increased methylation of archaeal tetraethers at elevated temperatures (Sollich et al., 2017). The most widely recognized adaptation, however, involves an increased proportion of tetraethers in membranes, as observed in *Thermococcus barophilus* and *Thermococcus kodakarensis* that both have a T_max_ = 100 °C (Cario et al., 2015; Matsuno et al., 2009) **(Table 1).**

**Table 1:** Comparison of the diether/ tetraether ratio for different archaeal species as a function of the optimum growth temperature. D= diether and T= tetraether. The dotted line separate mesophilic from hyperthermophilic species.

| Species | Order | Kingdom | Optimal growth temperature (°C) | Lipids |  | References |
| --- | --- | --- | --- | --- | --- | --- |
|  |  |  |  | D | T |  |
| *N. piranensis (C)* | Thaumarchaeota (Nitrosopumilales) | Proteoarchaeota | 25 | 8 | 68 | Elling *et al.*, 2017 |
| *M. boonei* | Stenosarchaea (Methanomicrobiales) | Euryarchaeota | 35 | 4 | 96 | S. Bräuer *et al.*, 2011 |
| *M. concilii* | Stenosarchaea (Methanosarcinales) | Euryarchaeota | 35 | 70 | ND | C.G. Choquet *et al.*,1992 |
| *T. acidophilum* | Thermoplasmatales | Euryarchaeota | 55-59 | 90 | 10 | N. Nemoto *et al.*, 2003 |
| *M. Prunae* | Crenarchaeota (Sulfolobales) | Proteoarchaeota | 75 | Traces | 70 | Itoh *et al.*, 2001 |
| *S. acidocaldarius* | Crenarchaeota (Sulfolobales) | Proteoarchaeota | 80 | 25 | 75 | Zeng *et al.*, 2018 |
| *T. kodakarensis* | Thermococcales | Euryarchaeota | 85 | 41 | 59 | Y. Matsuno *et al.*, 2009 |
| *T. barophilus MP* | Thermococcales | Euryarchaeota | 85 | 55,1 | 44,3 | M. Tourte *et al.*, 2021 |
| *A. pernix* | Desulfurococcales | Proteoarchaeota | 90-95 | 100 | 0 | J. Kejzar *et al.*,2024 |
| *I. aggregans* | Thermoproteota (Desulfurococcales) | Proteoarchaeota | 93,5 | Traces | 61 | Knappy *et al.*, 2011 |
| *M. kandleri* | Methanopyrales | Euryarchaeota | 98 | 100 | 0 | D. Hafenbradl *et al.*, 1996 |
| *P. furiosus* | Thermococcales | Euryarchaeota | 100 | 30 | 50 | M. Tourte *et al.*, 2021 |

Nevertheless, tetraether-containing membranes are not a prerequisite for heat tolerance **(Table 1).** For instance, *Methanopyrus kandleri* and *Aeropyrum pernix* exhibit little to no tetraethers, yet they grow optimally between 95 °C and 105 °C (Hafenbradl et al., 1996; Ulrih et al., 2009). Moreover, the deletion of the tetraether synthase (Tes) enzyme in *Thermococcus kodakarensis* does not prevent growth at 95 °C (Liman et al., 2024). Interestingly, tetraether lipids are also found in mesophilic Archaea, such as *Methanogenium boonei* (T_opt_ = 35 °C) and *Nitrosopumilus piranensis* (T_opt_ = 25 °C).

To elucidate these observations and assess the necessity of tetraethers for membrane stability at high temperatures, we investigated the behavior of archaeal-like bilayer membranes containing varying concentrations of tetraethers. Using purified lipids from *Pyrococcus furiosus* and *Sulfolobus acidocaldarius*, we conducted neutron diffraction experiments under high-temperature conditions to mimic the extreme environments inhabited by Archaea. This approach allowed us to characterize the ultrastructural response of the membranes to temperature variations as a function of tetraether concentration. It also provided crucial insights into how membrane architecture adapts to thermal stress, shedding light on the fundamental principles of cellular resilience. Understanding these mechanisms not only deepens our knowledge of archaeal survival strategies but also offers potential applications in biotechnology, such as engineering heat-resistant biomaterials and developing robust synthetic membranes for industrial processes.

## Materials and Methods

### Cell cultures

Diether lipids were purified from the hyperthermophilic archaeon *Pyrococcus furiosus* strain DSM3638 (Deutsche Sammlung von Mikroorganismen und Zellkulturen, DSMZ). Cultures were carried as described previously (Lipscomb et al., 2011) in 18 L of Defined Cellobiose (DC) minimal medium at 98 °C in a 20 L fermenter under strict anaerobiosis. The culture was inoculated to 1.10^6^ cells/mL and incubated at 95 °C for 20 h. Cells were recovered by centrifugation in 1 L batches, rinsed twice in sterile 3 % NaCl aqueous solution and lyophilized overnight.

Tetraethers were purified from the acidophilic and thermophilic archaeon *Sulfolobus acidocaldarius* strain MW001 (Wagner et al., 2012). Cells were grown aerobically at 75 °C with shaking in Brock’s minimal medium in the presence of 20 μg.mL^−1^ uracil (Sigma-Aldrich) (Brock et al., 1972). Cells were recovered by cross-flow filtration, rinsed twice, and lyophilized overnight.

### Diether lipid purification from *P. furiosus* cell pellets

Intact polar lipids (IPLs) from *P. furiosus* were extracted with 40 mL of a monophasic mixture of methanol (MeOH)/trichloromethane (TCM)/purified water (1:2.6:0.16; v/v/v) using a sonication probe for 10 min. After centrifugation (4000x rpm, 5 min), the supernatant was collected, and the extraction procedure was repeated twice. All extracts were pooled and dried under reduced pressure using a rotary evaporator. The dried residue was solubilized using a mixture of MeOH/TCM (1:1; v/v), transferred into a 2 mL vial, and the solvents removed under an N_2_ stream. Extracted intact polar lipids were kept at -20 °C until lipid purification.

Diethers were purified by 2D TLC as described by Christie (2011) on silica-gel pre-coated glass-backed plates (60 Å silica-gel, 20 cm x 20 cm plates, 0.5 mm SiO_2_ thickness). The plates were first developed in the first direction in the 1D mobile phase mixture TCM/MeOH/water (75:25:2.5; v/v/v). The plates were fully dried and developed at right angle in the second direction in the 2D mobile phase TCM/MeOH/acetic acid/water (80:9:12:2; v/v/v/v) (Christie, 2011). Lipids were detected with iodine and collected by scrapping the silica gel zone. The purified lipids were extracted from the silica gel with a mixture of TCM/MeOH (1:1; v/v) **(Figure 1)**. See supplementary for the detailed procedure.

**Figure 1:**
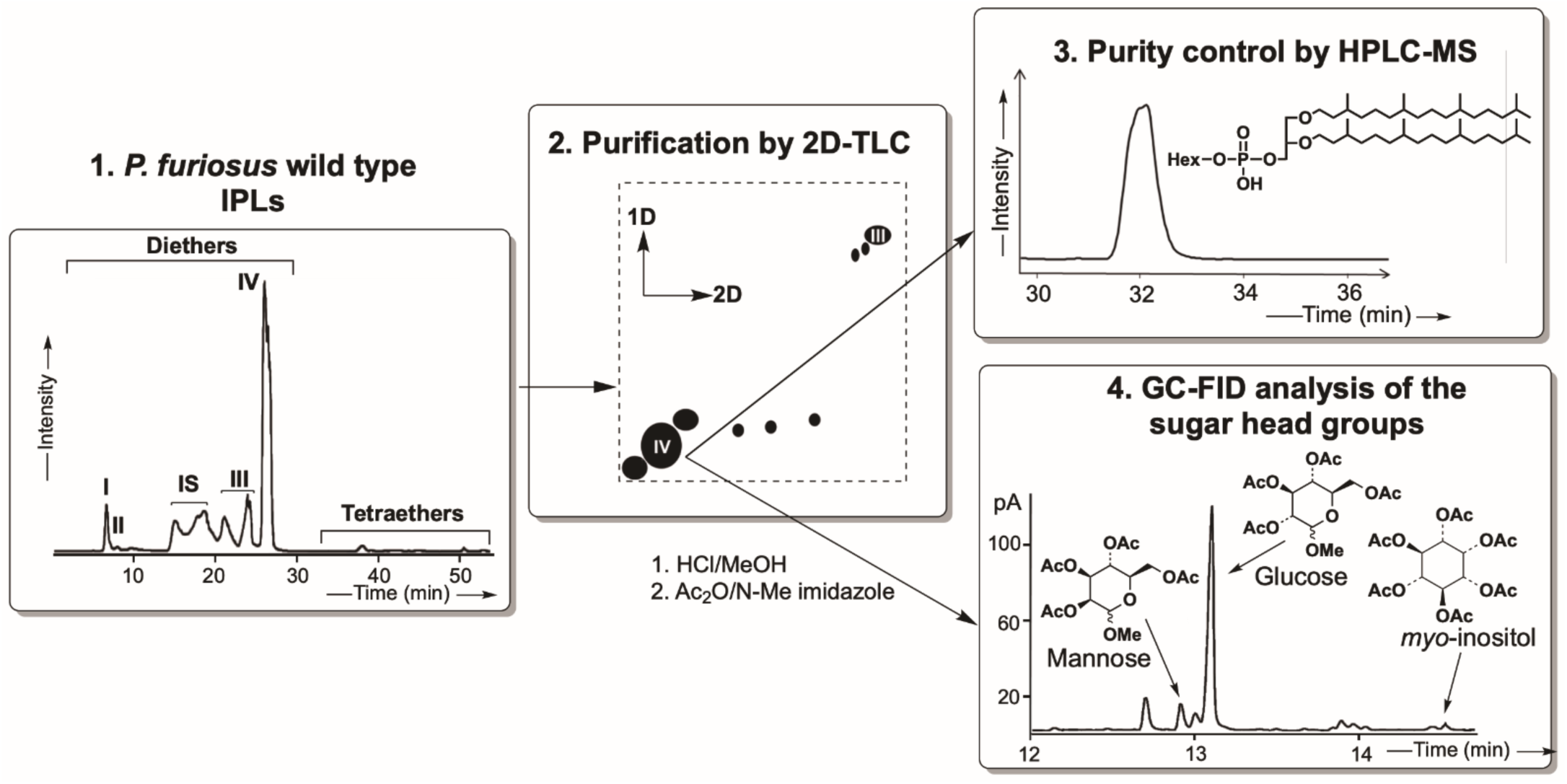
Schematic representation of PI-DGD purification steps and identification of the hexose head groups. **1.** HPLC-ESI-MS (negative ions) base peak chromatogram of IPLs of *P. furiosus* wild type strain; I: P-DGD; II: PG-DGD; III: PHexNHAc-DGD; IV: PHex-DGD. **2.** Purification of PHex-DGD by two-dimensional thin layer chromatography (1D= (TCM/MeOH/H_2_O; 75:25:2.5; v/v/v); 2D= (TCM/MeOH/AcOH/H_2_O; 80:9:12:2; v/v/v/v) (after Christie, 2011). **3.** HPLC-ESI-MS (negative ions) base peak chromatogram for controlling the purity of the isolated compound PHex-DGD. **4.** Methanolysis of the purified PHex-DGD followed by acetylation and GC-FID analysis to identify the hexose head groups.

The IPL content of each TLC extract was verified by HPLC-MS using an HP 1100 liquid chromatography module (Agilent) coupled to an HCT mass spectrometer (Bruker) equipped with an electrospray ionisation (ESI) source used in positive and negative ionisation mode. The parameters of the ion source were as follows: source temperature 350 °C; capillary transfer voltage +5000 V (negative mode) and - 4000 V (positive mode); nebulizer pressure 43 psi; drying gas flow rate (N_2_) 8 min^-1^; drying temperature 350 °C; corona discharge 4 μA. The detection range (m/z ratio) was 500 to 3000 m/z. N_2_ was generated from pressurised air by a Calypso 25.0 (F-DGSi) nitrogen generator. Chromatographic separations were carried out using an Inertsil Diol HILIC column (2.1 x 250 mm, 5 µm; GL Science) thermostated at 30 °C and equipped with a precolumn having the same stationary phase. The separation was carried out in solvent gradient mode (modified after Jaap S. Sinninghe Damstï et al., 2012) and included as the mobile phase solvent A (isopropanol (IPA)/water/formic acid/aqueous ammonia (88:10:0.12:0.04 v/v/v/v) and solvent B (*n*-heptane (*n*-C_7_)/IPA/formic acid/aqueous ammonia (79:20:0.12:0.04 v/v/v/v)). Lipids were eluted at a constant flow rate of 0.2 mL min^-1^ in a stepwise gradient from 100 % solvent B to 66 % solvent B over 18 min, then maintained for 12 min, followed by 35 % solvent B in 15 min, maintained for 15 min and then back to 100 % solvent B in 2 min, followed by stabilisation of the HPLC column for 20 min. The IPL were solubilised in 1 mL of solvent B. Injections of 10 µL were performed using an Agilent autosampler. Analyses were processed using Data Analysis software (version 4.2, Bruker Daltonics).

### Tetraether purification from *Sulfolobus acidocaldarius* cell pellet

*Sulfolobus acidocaldarius* cell pellets were provided by M. Tourte and S.V. Albers from the Molecular Biology of Archaea team (Institute of Biology II, University of Freiburg, Germany). Most previous studies on tetraether lipids have been performed on the polar lipid fraction E (PLFE) of *S. acidocaldarius*. Our preliminary HPLC-MS analyses of this PLFE fraction showed that this fraction was composed of several different tetraether lipids. In order to better interpret the results of the current experiment, it was necessary to try to further purify this tetraether mixture. The complete purification procedure involved a protection/deprotection sequence and IPL purification by HPLC, and will be described in detail elsewhere (Schaeffer et al. in preparation).

### Core lipid characterization

Core lipids from *P. furiosus* and *S. acidocaldarius* were obtained as described previously (Tourte et al., 2020a). Briefly, aliquots of the isolated di-or tetraether polar lipids were hydrolyzed in Pyrex^TM^ tubes with PTFE caps by acid methanolysis (1.2 N HCl in MeOH).For the analysis of the core lipids from *P. furiosus*, a Zorbax Sil column (4.6 x 250 mm, 5 μm; Agilent) was used in isocratic mode with a pre-column having the same stationary phase. The mobile phase consisted of a *n*-C_7_/IPA solvent mixture (95:5 v/v) at a flow rate of 0.4 mL min^-1^. The analysis time was set at 40 min, the core lipids were solubilized in n-C_7_/IPA (99:1 v/v) and the injection volume was set at 10 μL.

For the analysis of the core lipids from *S. acidocaldarius*, a Zorbax Rx-Sil column (2.1 x 150 mm, 5μm; Agilent) with a pre-column having the same stationary phase was used in solvent gradient mode with solvent A (n-C_7_/IPA 98.5:1.5 v/v) and solvent B (IPA) being used with the following gradient: 100 % solvent A for 10 min, then 2 % solvent B in 20 min, then increase to 10 % solvent B in 30 min followed by 14 % solvent B in 55 min, back to 0 % solvent B at 56 min and 19 min equilibrium period of the HPLC column with 0 % solvent B. Lipids were eluted at a constant flow rate of 0.2 mL min^-1^.

The following MS parameters were applied for the analysis of the core lipids from both *P. furiosus* and *S. acidocaldarius*: Capillary voltage: -2000 V, corona +4000 nA, nebulization pressure 50 psi, drying gas flow rate (N_2_) 5 L min^-1^, drying temperature 350 °C, vaporization temperature 420 °C, mass scan from 500 to 3000 m/z.

### Headgroups characterization

The polar crude mixture obtained upon acidic hydrolysis of isolated polar lipids (i.e., the part that was not dissolved in n-C7/IPA 99:1 v/v) was transferred into a 3 ml vial using a DCM/MeOH (1:1 v/v) mixture, and the solvent was removed under a stream of argon. After addition of ethyl acetate (EtOAc) (ca. 1 mL), the crude mixture was acetylated using 200 µl of Ac_2_O and 100 μl of N-Me-imidazole. After 1 h at room temperature, ca. 2 mL of distilled water containing 25% CuSO_4_ (v/w) were added, and the mixture was vigorously shacked. The supernatant (i.e., the EtOAc phase) containing the acetylated methoxy-sugars was recovered using a Pasteur pipette and analysed by GC-FID.

GC-FID analyses of the acetylated methoxy sugars released by acid hydrolysis (see above) were carried out on a Hewlett Packard 6890 gas chromatograph equipped with an on-column injector used in the “track-oven” mode, a flame ionization detector, and an HP-5 fused silica capillary column (30 m x 0.25 mm; 0.25 μm film thickness; Agilent). H_2_ was used as carrier gas (constant flow mode, 2.5 mL min^-1^), and the oven was programmed as follows: 70–200 °C (4 °C min^-1^), 200–300 °C (10 °C min^-1^), isothermal at 300 °C. The different acetylated methoxy sugars formed upon methanolysis/acetylation were identified by comparing their retention times in GC with those from reference compounds.

### Archaeal bilayer membrane reconstructions

To assay the relative contribution of each type of lipid (diether vs. tetraether) on membrane features and properties, we synthesized 4 types of samples with various diether to tetraether (D/T) ratios. In addition to the purified diether (6 mg) and tetraether (3 mg), which served as pure pole controls, were prepared 2 D/T mixtures: a 2:1 (molar) D/T mixture containing 3 mg of diether and 3 mg of tetraether, and a 1:1 (molar) D/T mixture containing 1.5 mg of diether and 3 mg of tetraether. The four samples were studied as a multi-stack of lipid bilayers on one-sided polished ultraclean silicon wafers with a thickness of 275 ± 25 μm purchased from Si-Mat (Germany). The wafers were previously cut to produce a rectangular shape (4 cm x 2.5 cm) to fit the sample compartment. Silicon wafers were rinsed with TCM and dried by a N_2_. Lipid samples were spread on a silicon wafer and dried overnight under vacuum.

Samples were transferred to the high-precision BerILL humidity chamber (Gonthier et al., 2019) and mounted vertically on a manual 4-axis goniometer head (Huber, Germany) set in the humidity chamber and pre-aligned using a laser-based optical setup. For the four samples, at least 3 temperatures (60 °C, 70 °C, and 80 °C), and 2 relative humidities (80 % and 95 % RH) were tested. Each sample was incubated in the humidity chamber for at least 3 hours prior the first data collection. Among the different conditions, 1 h of equilibrium was respected to ensure a constant *d*-spacing and constant intensity of Bragg reflections during the experiment.

### Neutron diffraction: d-spacing calculation

Neutron diffraction data were collected at the recently upgraded D16 cold neutron diffractometer of the Institut Laue-Langevin (ILL, Grenoble, France) (Cristiglio et al., 2015). The instrument design has been revisited, with a new secondary spectrometer to allow the installation of a new curved wide-angle 2D-detector, based on the trench technology developed at ILL (Buffet et al., 2023). The new detector has a pixel resolution of 1.5 × 2.0 mm^2^ (hor. X vert.) providing an angular resolution of 0.075 x 0.1 deg. 211q resolution at a distance of 1150 mm.

To collect the data, the vertical sample plane is illuminated by the horizontally collimated and vertically focused incident beam with an adjustable angle of incidence set by the sample angle Ω. The beam is scattered in different directions at angles 2θ from the incident beam. For each Ω, the 2θ dependent intensity is collected. By rotating the sample stage, Ω-scans (rocking curves) were performed by steps of 0.05 deg. from -1 to 13 deg. The intensity was corrected for the detector pixel efficiency resulting from a water calibration run. For each Ω-scan, 2D images obtained at a given Ω step were reduced to 1D by vertical integration of the intensity in an ROI (2θ_y_ *vs.* 2θ_x_ range) centered vertically on Bragg reflections, and stacked to produce the reciprocal space map. These steps from the Ω-scans’ raw data to the production of the 2D reciprocal space maps were performed using the Mantid software, Daily version (Arnold et al., 2014). The analysis of the reciprocal space maps was done in Igor Pro 8.0 (WaveMetrics, Lake Oswego, OR, USA) using a dedicated procedure written by T. Hauss for the new D16 detector and data format.

The analysis consisted in the integration of the intensity along the specular direction to produce the I vs 2θ curve. Bragg peaks were then fitted using the Multipeak fitting tool of Igor Pro, the positions and intensities of which were used for the determination of the period of the lamellar membrane stack (d-spacing) and for the reconstruction of the membrane density profile in real space. Here, q_z_ is the scattering vector normal to the bilayer planes related to the scattering angle (2θ) defined as:

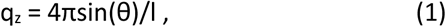

When several orders of diffraction were observed (up to 4), a linear fit of the peak positions vs. Bragg order (n) was performed. The slope of which was used to determine the d-spacing (d) using the equation (1), where λ is the incident neutron beam wavelength (4.487 Å): d = λ/Δq. When only a single Bragg reflection was observed, the d-spacing was calculated using the first-order peak according to d = 2p/q_z_. The error on the d-spacing was given by propagating the error on the slope of the linear fit to the data.

### Neutron scattering length density (NLSD) profiles: membrane bilayer and water layer thicknesses

NSLD profiles were calculated according to (Harroun et al., 2008) from the integrated intensities of Bragg peaks corrected for the neutron absorption (C_abs_), the Lorentz correction (C_Lor_), and the neutron flux correction (C_flux_), resulting in the corrected discrete structure factor of order n:

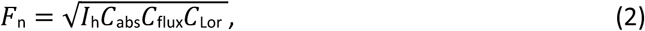

where I_h_ is the intensity of the Bragg peak at the order *h.* The corrections are given by:

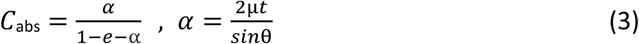

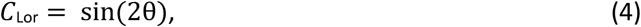

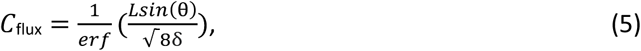

where *t* is the sample thickness calculated from the deposited amount of dry lipid and the sample area, 2δ is the beam width (Hrubovčák et al., 2019).

Finally, the NSLD was calculated using:

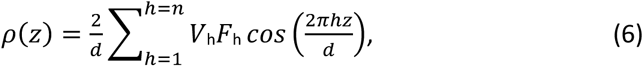

where z is the specular direction perpendicular to the bilayer planes, *h* the diffraction order, *n* the highest order detected and V*_h_* corresponds to the phase of the structure factor *h*. Using the 8 % D_2_O contrast, lipid headgroups are highlighted due to the zero NSLD of water and the negative NSLD of aliphatic chains. We tested the different hypotheses for the discrete structure factor signs to obtain an NSLD profile with a minimum at z = 0 and a maximum NSLD for the polar head regions. The best agreement with these constraints was [-, -, +, -] for V_h_. From these profiles, the Gibbs–Luzzati bilayer thickness (*d_b_*) can be directly extracted (Luzzati and Husson, 1962), as the center-to-center distance between polar headgroup layers (Salvador-Castell et al., 2020). The thickness of the water layer between the lipid bilayer (*d_w_*) is calculated from the known *d*-spacing and the bilayer thickness according to: d_w_ = d - d_b_, as illustrated **(Figure S1 and Table S1).**

## Results and discussion

### Purified lipid characterization

To better mimick the biological reality, in contrast to previous studies that used synthetic, archaeal-like lipids, we chose to perform our measurements on natural, isolated polar lipids rather than on lipid mixtures, synthetic lipids, or homologs.

The diether lipid (D) used in our experiment consisted of a dialkylglycerol diether (DGD) with a phosphatidyl hexose polar headgroup (PHex-DGD; [M-H]^-^ 893, **Figure 1**). Following extraction and purification from *P. furiosus* lipid extract, the acid hydrolysis followed by acetylation of an aliquot of the isolated PHex-DGD revealed a polar headgroup composed of a mixture of methoxy sugars, comprising glucose (Glc) (ca. 89 %), mannose (ca. 9 %) and inositol (ca. 2 %). However, we could not separate these different diethers given that the different hexose polar headgroups have the same polarity on silica gel and thus cannot be separated by the 2D TLC method used and were also not separated by the HPLC-MS analytical conditions used. These stereoisomers or derivative structures elute at the same time and have the same mass (Shen and Perreault, 1998; Zuo et al., 2020). Although this isolated fraction did not consist of a single compound, the polar head group composition was largely dominated by glucose, and for this study, we assumed that the diether we worked with was PGlc-DGD **(Figure 1)**.

Regarding the tetraether lipids (T), we decided to work on the famous archaeon *S. acidocaldarius* polar lipid fraction E (Lo and Chang, 1990). However, our preliminary analyses showed that this fraction was a too complex lipid mixture. To obtain pure tetraethers, we designed a novel approach to separate the different tetraether lipids within this fraction, which involved the protection and deprotection of lipids (Schaeffer et al, in preparation). After several attempts, we could reduce the number of lipid molecules in the extracts but could not get samples containing only a single lipid molecule. The best that we could obtain was a mixture of two different glycerol dialkylglycerol tetraether series comprising P-Hex2-GDGT-0-6 ([M-H]^-^ 1705-1693 Da, **Figure 2A 2C**), and P-Hex3-GDGT-0-6 ([M-H]^-^ 1867-1854 Da, **Figure 2A 2C**). For both series, a combination of tetraethers having two C40 isoprenoid carbon (biphytane) chains with 0 to 6 cyclopentane rings was present **(Figure 2C)**. Last, acid hydrolysis followed by acetylation of this tetraether mixture revealed that the hexose groups were mostly glucose (ca. 98 %) rather than inositol (ca. 2 %) **(Figure 2B)**.

**Figure 2:**
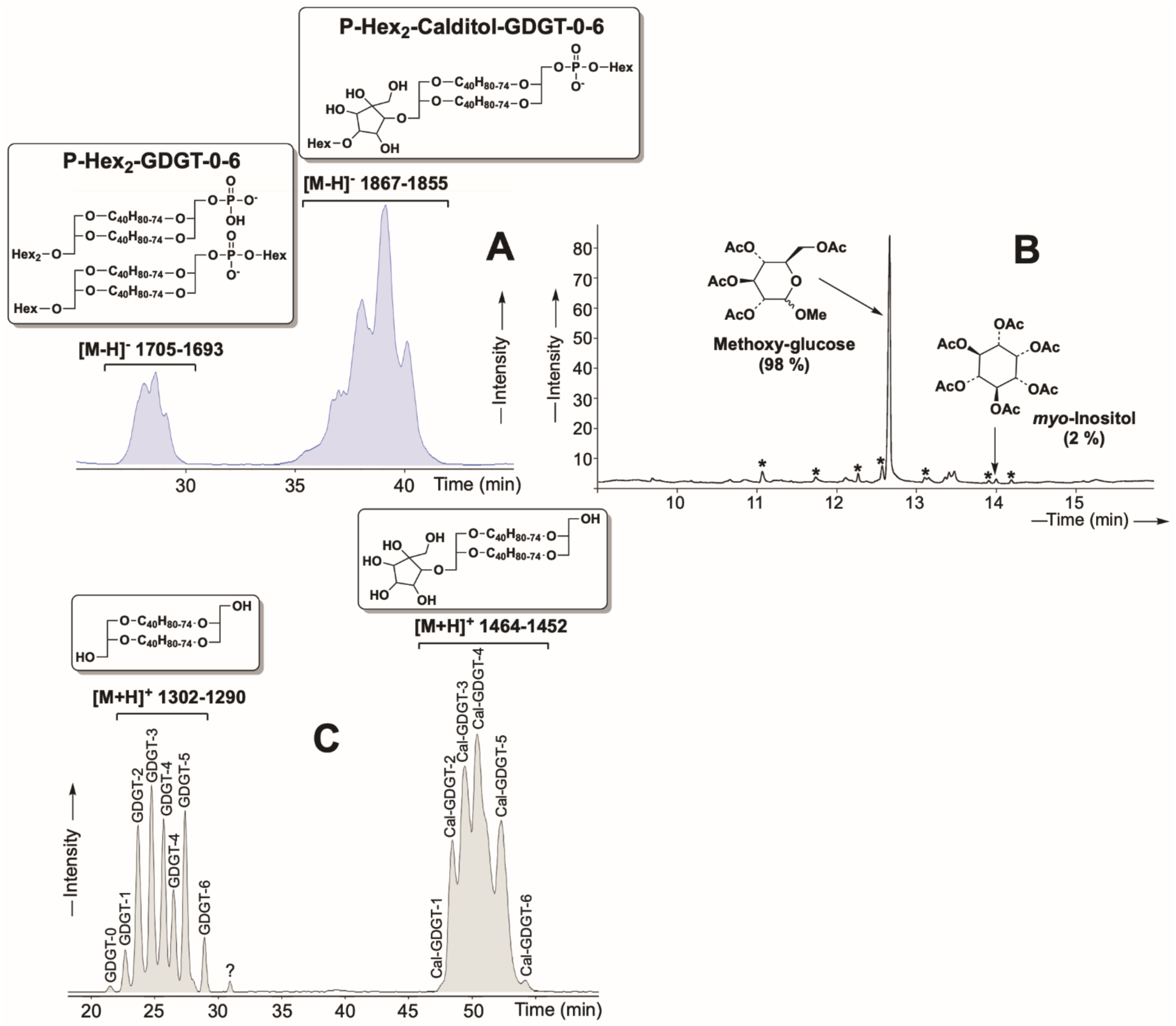
Characterisation of the tetraether mixture isolated from the lipid extract from *S. acidocaldarius*. **A.** HPLC-ESI-MS (negative ions) base peak chromatogram of the tetraether polar lipid fraction isolated *from S. acidocaldarius*. **B.** Gas chromatogram (GC-FID) showing the sugar head groups recovered upon methanolysis of the isolated tetraether mixture from *S. acidocaldarius*. The sugars are analysed as acetate derivatives. * : contaminations **C.** HPLC-APCI-MS (positive ions) base peak chromatogram of the core lipids recovered upon methanolysis of the tetraether polar lipid fraction isolated from *S. acidocaldarius*. GDGT-0-6: Glycerol dialkyl glycerol tetraether with 0 to 6 cyclopentane rings. Cal-GDGT-0-6: Calditol-GDGT with 0 to 6 cyclopentane rings.

### The importance of natural polar headgroups on membrane stability

The ability of our natural lipids to form oriented bilayer structures was confirmed using neutron diffraction on the D16 instrument at the ILL. Typically, lipid films hydrated by water vapor form stacks of bilayers separated by water layers. The multilayer nature of the sample leads to the appearance of Bragg peaks in the diffraction data. In our study, the stacked multilayers were sufficiently ordered to produce Bragg peaks across all tested conditions for our four samples **(Figure 3A)**. Notably, strong signals were obtained from our natural lipid sample even at elevated temperatures of 80 °C or 90 °C, which was the high-temperature limit of the BerILL setup. This is in contrast to synthetic archaeal-like diethers (DoPhPC, or DoPhPE), which displayed poorly structured bilayers at comparable temperatures (LoRicco et al., 2020; Salvador-Castell et al., 2021, 2020), which validates the use of natural vs. synthetic lipids for the exploration of archaeal membrane biophysical properties.

**Figure 3:**
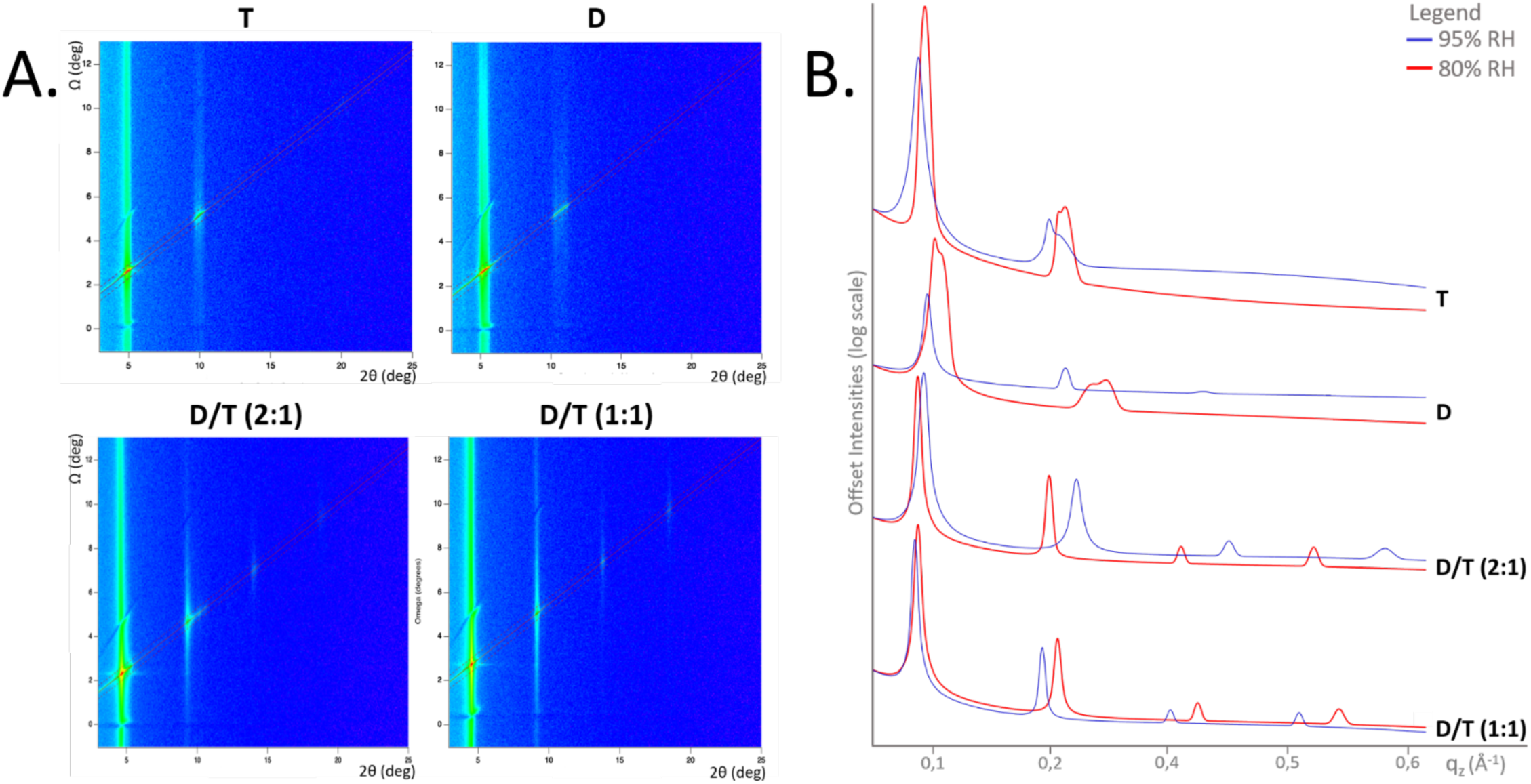
Neutron diffraction of multilayer samples with varying concentrations of purified tetraether (T) and diether lipids (D). **A.** Reciprocal space maps resulting from typical Ω-scans*I* (2θ, Ω) showing the Bragg peak positions of the different lipid samples, run at 100 D_2_O 70 °C 80 RH. **B.** 1D integrated intensity along the Z direction of the different lipid samples highlighting Bragg peak intensities, run at 100 D_2_O 70 °C. Red lines represent 80 % RH measurements, and blue lines 95 % RH measurements. D= diether sample, T=tetraether sample, D/T=mixture of diether and tetraether at different molar ratios (1:1) or (2:1).

Hence, the enhanced stability of our reconstructed membranes appeared to be closely linked to the polar headgroup composition of the natural lipids used. Previous studies have been performed with synthetic archaeal-like lipids bearing headgroups such as phosphatidylethanolamine (PE) or phosphatidylcholine (PC), which are common bacterial phospholipid headgroups, but are rarely seen in Archaea. Indeed, PE is prevalent in *E. coli* membrane lipids (Sohlenkamp and Geiger, 2016) but less dominant in Archaea, where it has been detected in the membrane lipids of only a few archaeal species to date, as *Methanosarcina barkeri, Archaeoglobus fulgidus,* and *Thermococcus kodakarensis* (Meador et al., 2014; Nishihara and Koga, 1995; Tarui et al., 2007). PC, common in eukaryotic cells, is even rarer in archaeal membranes, with reports limited to *Methanopyrus kandleri* (Nishihara et al., 2002). Archaeal membranes have been shown to typically feature phosphatidylinositol (PI) or phosphatidylglycerol (PG) or more complex structures such as oligosides, sulfur-substituted sugars, or a methylphosphate PG. Synthetic analogs of these lipids are scarce due to the complexity of their synthesis and are not commercially available. Several studies have shown that polar headgroup size and structure critically influence membrane packing and stability. For instance, the smaller PE headgroup results in tighter membrane packing, whereas the larger PC headgroup produces a less densely packed membrane (Kodama and Miyata, 1996; Winter and Jeworrek, 2009). In the specific case of archaeal-like lipids, a 9:1 DoPhPC:DoPhPE mixture maintained organized membrane structures up to 70 °C. Interestingly, archaeol-based lipids with PI headgroups retained lamellar organization only between 25 °C and 55 °C, deteriorating at higher temperatures (Ruiz et al., 2023). In contrast, our PHex-DGD diether sample, harboring essentially glucose (89 %) and mannose (9 %) polar headgroups, demonstrated enhanced stability, producing three Bragg peaks at 60 °C and 70 °C at 95 % relative humidity. This suggests that the nature of the polar headgroups of hyperthermophilic strains is an essential component of the membrane physico-chemical properties, and participates strongly in archaeal bilayer membrane auto-organization and stability. In the example of the extremophilic archaeon *P. furiosus*, from which these lipids were purified, we hypothesize that this may be due to the distinct properties of phosphoglucose and phosphomannose vs. PC or PE, or to the mixed composition of our PHex-DGD extract **(Figure 1)**. In fact, compared to the PC and PE headgroups, sugars can create an H-bonding network with water. Intra vs intermembrane interactions bridging results in an attractive contribution between membranes that stabilizes the lamellar phase (Kanduč et al., 2017).

Our experiment performed on purified lipids is congruent with previous observations made on archaeol lipid mixtures. For example, in the acidophile *Picrophilus oshimae*, the intact polar lipids extracted are incapable to self-organize into liposomes at pH above 4, while they form extremely stable and impermeable liposomes at lower pH (van de Vossenberg et al., 1998). Similarly, purified lipids extracted from extreme halophiles like *Halobacterium halobium* do not form liposomes in the absence of NaCl in the buffer, while at high NaCl concentrations, liposomes exhibit extreme membrane stability, which has been attributed to the halophile-specific polar headgroup, phosphatidylglycerophosphate methyl ester (PGP-Me). This headgroup is predicted to prevent bilayer aggregation through steric repulsion (Tenchov et al., 2006). These findings highlight how the polar headgroups of natural lipids have evolved to mimic/counteract the environmental conditions in extremophilic Archaea.

### Lipid diversity as a key to membrane stability and adaptability

One of the questions raised for membranes composed of a mixture of bilayer and monolayer-forming lipids is how these organize spatially and how they interact/cooperate (or not) to form membranes. Our structural analyses revealed that membranes containing a mixture of diethers and tetraethers (D/T mixtures) were more ordered than those composed solely of one lipid type. Ω-scans and 1D neutron diffractograms showed that D/T mixtures exhibited 4 well-defined Bragg peaks, indicating a well-organized structure compared to the broader peaks observed in pure diether and tetraether samples **(Figure 3AB)**. At 80 % RH, the broad scattering patterns for T and D membranes suggested inhomogeneities due to two coexisting lamellar phases, consistent with previous studies (Salvador-Castell et al., 2020) **(Figure 3A, 3B)**. At 95 % RH, the diether mixture D rearranged into a unique lamellar phase with three Bragg orders, while the tetraether film remained inhomogeneous **(Figure 3B)**. D/T mixtures also exhibited consistent d-spacing values across tested temperatures and humidity around 55 Å whereas pure tetraether membranes showed greater variability **(Figure 4)**. These results underscore the role of lipid diversity in maintaining membrane integrity, suggesting that mixed lipid membranes are better equipped to adapt to environmental stresses. This may appear counterintuitive given the difference in molecule length between one tetraether molecule and two diether molecules.

**Figure 4:**
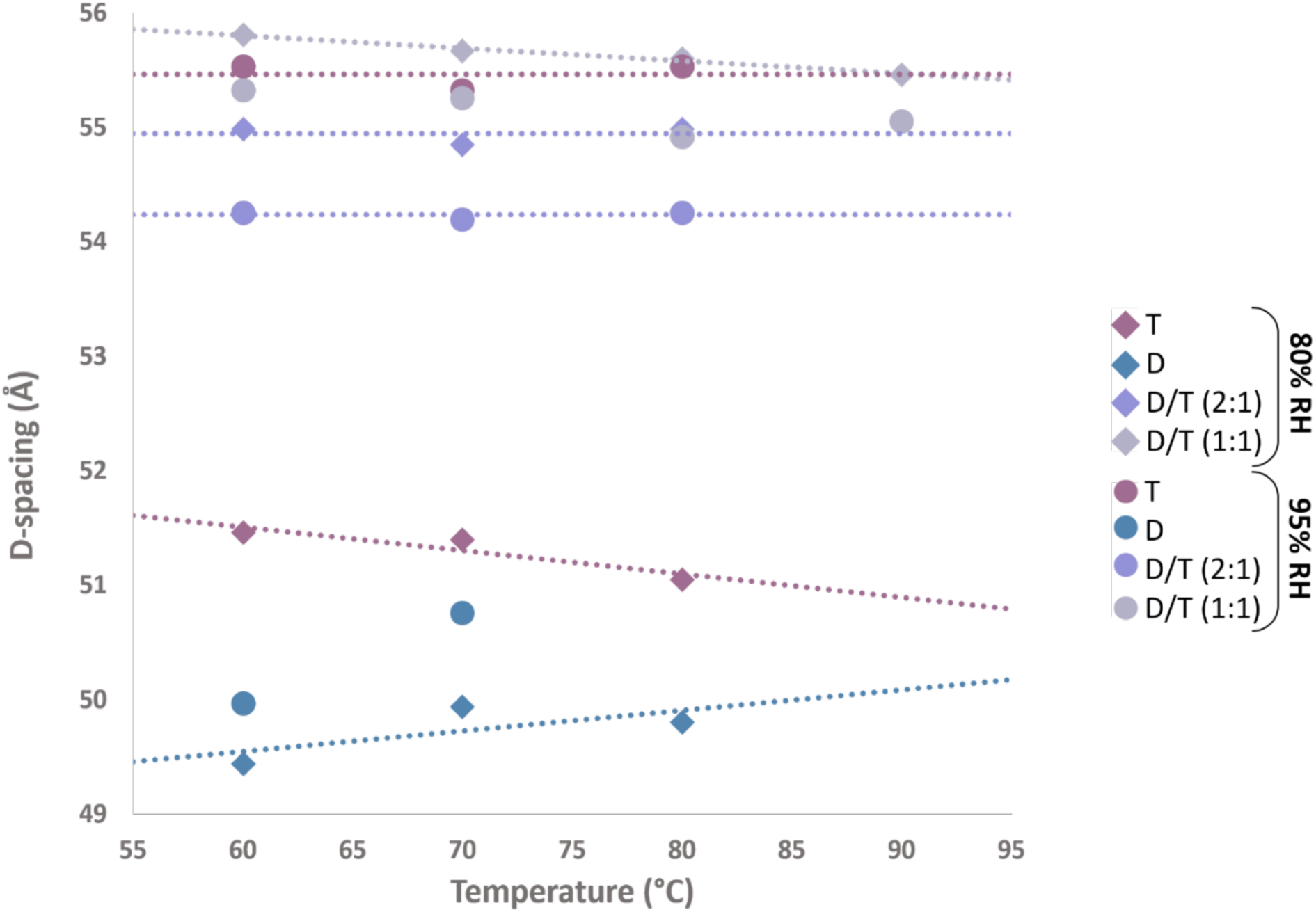
Lamellar d-spacing as a function of temperature and relative humidity for different lipid compositions. Measured at 80 % and 95 % RH at 60 °C, 70 °C and 80 °C. Only one sample was measured at 90 °C. The d-spacing was calculated only from 8 % D_2_O measurements (see methods). D= diether sample, T=tetraether sample, D/T=mixture of diether and tetraether polar lipids at different molar ratios (1:1) and (2:1). A problem occurred during the diether measurement at 80 °C 95 %RH, and it was thus not possible to calculate d-spacing for this sample.

NSLD profiles provided additional insights, calculated for all samples under six tested conditions with 8 % D_2_O. The quality of the NSLD profiles depends on the number and intensity of Bragg peaks. Compared to diether membranes, tetraether-rich membranes demonstrated higher structural resolution with more intense Bragg peaks **(Figures 5 and 6).** At 95 % RH, NSLD profiles showed for all samples the characteristic structure of a lipid bilayer, with a prominent peak corresponding to the phospholipid headgroup region and a central hydrophobic region **(Figure 6)**.

**Figure 5:**
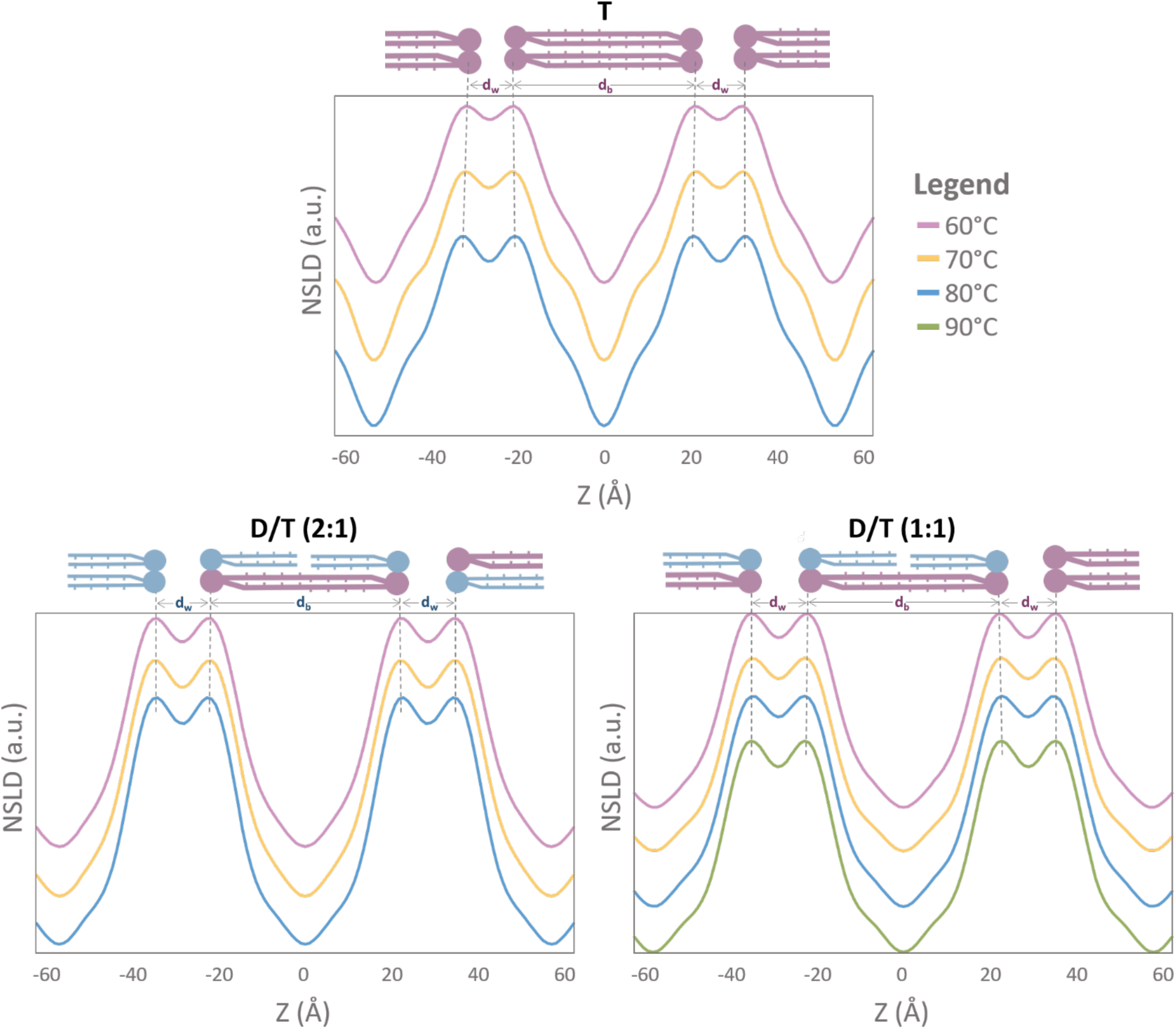
Neutron Scattering Length Density (NSLD) profiles showing two periods of different archaeal membrane compositions. Each plot corresponds to the NSLD profile at one humidity level (80 % RH) and different temperatures. The bilayer thickness d_b_ corresponds to the center-to-center distance between headgroups as described in the methods. The water layer thickness d_w_ is calculated according to d = d_w_ - d_b_, d being the d-spacing. The dotted black lines show the shift of the maxima in the NSLD profiles. **NSLD profiles were calculated with 4 for T, and both D/T samples.** T=tetraether sample, D/T=mixture of diether and tetraether samples at different molar ratios (1:1) or (2:1). In these conditions only 2 orders was detected for the diether sample (D), the NSLD profile was not calculated. Due to the lack of resolution normally provided by higher diffraction orders, diether membrane would lead to a wrong or featureless profile.

**Figure 6:**
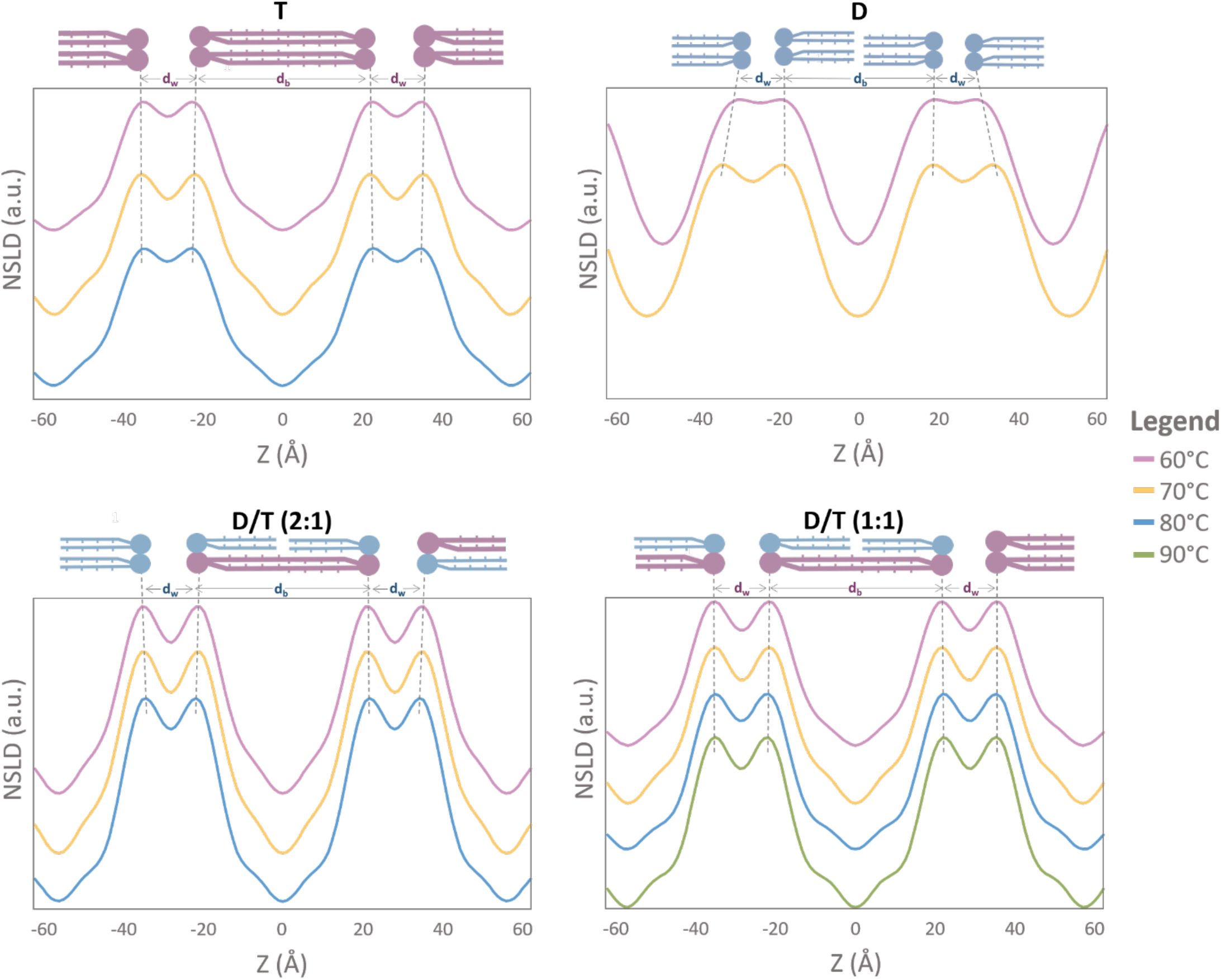
NSLD profiles showing two periods of different archaeal membrane compositions. Each plot corresponds to the NSLD profile at one humidity level (95 % RH) and different temperatures. The bilayer thickness d_b_ corresponds to the center-to-center distance between headgroups as described in the methods. The water layer thickness d_w_ is calculated according to d = d_w_ - d_b_, d being the d-spacing. The dotted black lines show the shift of the maxima in the NSLD profiles. **NSLD profiles were calculated with 3 orders for D samples and 4 for T, and both D/T samples.** D= diether sample, T=tetraether sample, D/T=mixture of diether and tetraether sample at different molar ratios (1:1) or (2:1). A problem occurred during the diether measurement at 80 °C 95 % RH, for this reason it was not possible to calculated NLSD profile for this sample.

Interestingly, temperature variations primarily affected the polar headgroup region, at -20 Å; +20 Å **(Figure 7)**. Increasing the temperature decreases the scattering signal. A warmer environment leads to disordered membranes, leading to less intense Bragg peaks. As this membrane region has been related to interactions existing within the lamellar phase, it appears that temperature variations cause reorganization over the polar headgroups. Hydration, however, had a more pronounced effect on hydrocarbon chain organization. For our lipid mixtures, the neutron scattering signal appeared less intense at 95 % RH compared to 80 % RH. Moreover, at 95 % RH, a more pronounced hump was observed in the hydrocarbon chain region for the tetraethers-containing membranes **(Figure 7)**. This hump visible in a region of 10 Å on each side of the bilayer midplane indicated different organization of branched chains in the lipid midplane and changes in hydrocarbon conformation **(Figure 7)**. This region is also associated with higher SLD carbonyl and phosphate groups (Kučerka et al., 2008; Shinoda et al., 2005). This modification is more noticeable for the tetraether sample T, meaning that the hydrocarbon chain is undergoing conformational changes while the humidity increases **(Figure 7)**. These findings suggested that the structural adaptability of tetraether membranes, characterized by flexible hydrocarbon chains, plays a crucial role in maintaining membrane stability under high humidity conditions.

**Figure 7:**
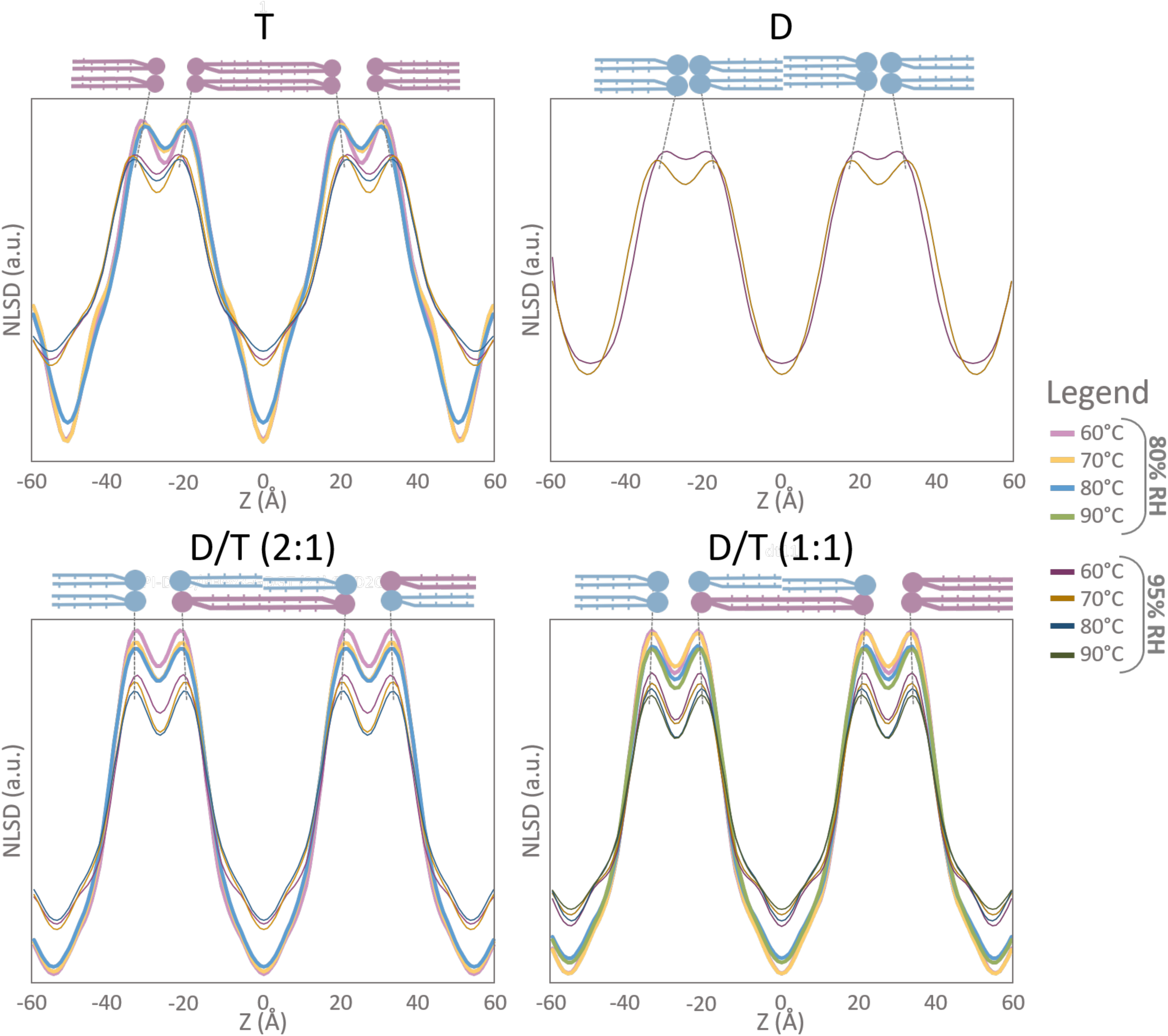
Overlaid NSLD profiles. Each graph corresponds to a NSLD profile at different temperatures and humidities. The dotted black lines show the shift of the maxima in the NSLD profiles**. NSLD profiles were calculated with 2 (80 % RH) or 3 (95 % RH) orders for D samples and 4 orders for T, and both D/T samples.** D= diether sample, T=tetraether sample, D/T=mixture of diether and tetraether sample at different molar ratios (1:1) or (2:1). At 80 % RH only 2 orders was detected for the diether sample (D), the NSLD profile was not calculated. Due to the lack of resolution normally provided by higher diffraction orders, diether membrane would lead to a wrong or featureless profile. A problem occurred during the diether measurement at 80 °C 95 % RH, for this reason it was not possible to calculated NLSD profile for this sample.

Membrane thickness measurements further underscored the stabilizing effect of lipid diversity **(Figure S1)**. The peak-to-peak distance in the membrane NSLD profile reflecting the center-to-center distance between polar headgroup layers (d-spacing), the Gibbs–Luzzati bilayer thickness (d_b_), and the thickness of the water layer between the lipid bilayer (*d_w_*) were calculated **(Figure 8A, 8B, Table S1)**. The measured membrane thickness of the diether samples decreased slightly when the temperature increased, from 39 Å at 60 °C to 36 Å at 70 °C, while the water thickness increased from 10 Å at 60 °C to 14 Å at 70 °C. Our results are congruent with previous studies on archaeal-like ether lipids which revealed a bilayer thickness of 38.3 Å at 25 °C and 40 °C (Salvador-Castell et al., 2020). No clear linear tendency was visible for T samples when comparing membrane thickness vs. temperature **(Figure 8B)**. However, at 80 °C, the calculated d_b_ increased by 3 Å, and the water thickness increased by 2 Å when increasing the humidity **(Figure 8A, 8B)**. The d-spacing variation observed for the T membrane is a combination of membrane thickness and water thickness variations. It can be due to changes in the degree of hydration of the polar headgroups, a tilt in the angle of the lipid molecules, or changes in the lipid conformation. In contrast, adding diether molecules to the membrane keeps all parameters stable **(Figure 8A, 8B)**. These results indicated that the combination of rigid and flexible lipid components in mixed membranes offers a balance that optimizes stability and adaptability, which appears to be crucial for survival in highly fluctuating environments that extreme environments are. These observations shed new light on the temperature/salinity/pressure adaptation response observed in the archaeal order Thermococcales, for which the homeoviscous adaptation has been shown to systematically involve a modification of the diether and tetraether lipid composition (Cario et al., 2015; Matsuno et al., 2009; Tourte et al., 2022, 2020b). In these species, which are capable of growth at 100 °C, the presence of diether lipids in the membrane may play an unanticipated and crucial role in their ability to adapt to highly fluctuating temperatures, as they thrive between the hydrothermal fluid, which temperature usually is above 350 °C, and the deep-sea, which temperature is in the range of a few degrees.

**Figure 8:**
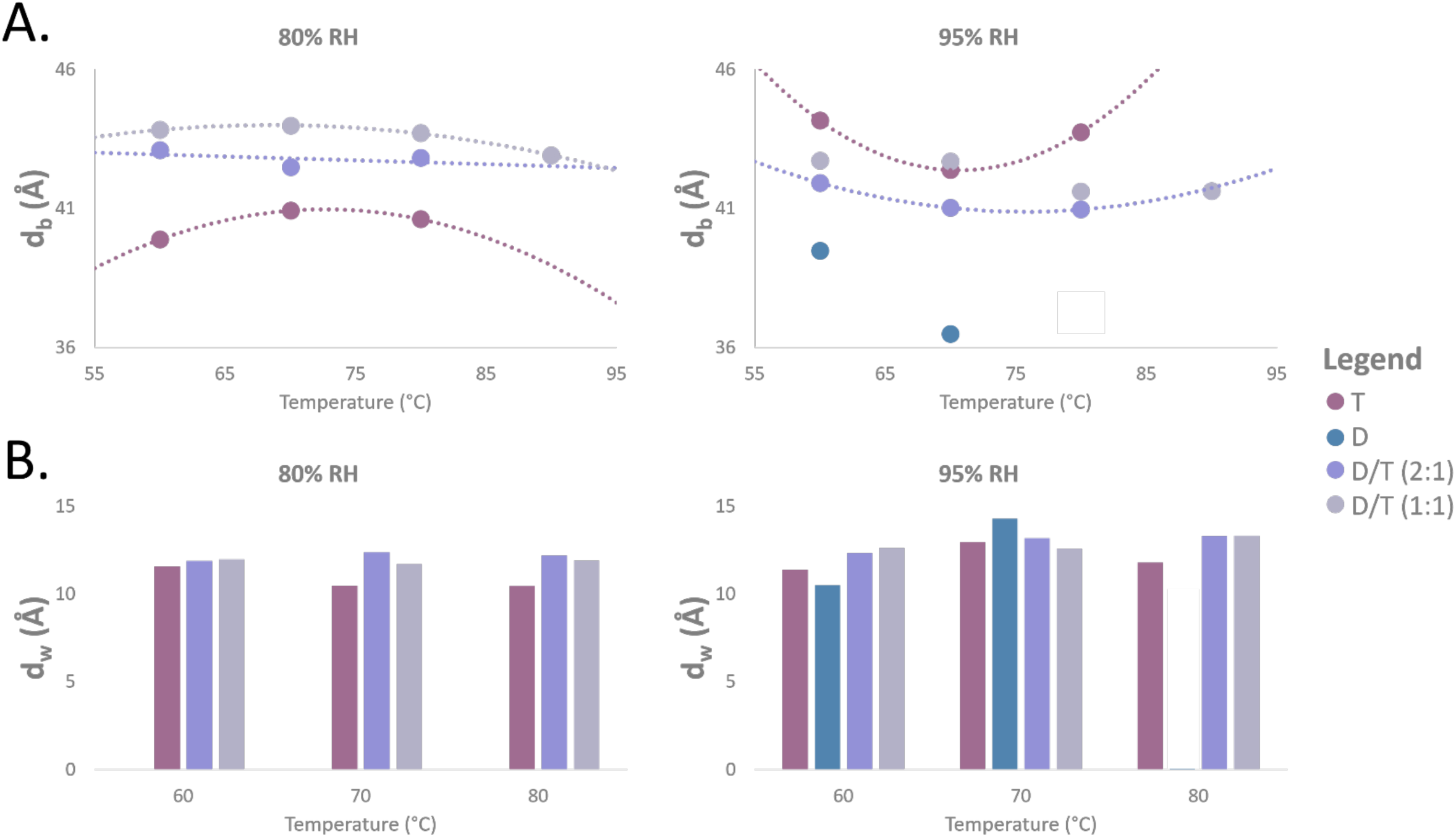
Thickness measurements for the different samples. **A.** Membrane thickness (d_b_) as a function of temperature for different lipids ratios, measured at 80 % and 95 % RH. Error for d_b_ measurements is ± 0.5 Å. **B.** Water thickness (d_w_) as a function of temperature for different lipids ratios, measured at 80 % and 95 % RH. Error for d_w_ measurements is ± 0.5 Å. D= diether sample, T=tetraether sample, D/T=mixture of diether and tetraether sample at different molar ratios (1:1) or (2:1). No data was available for the diether sample at 80 % RH for all tested temperatures, and at 80 °C with 95 % RH.

### Structural features of monolayer membranes

Tetraether membranes exhibit unique structural characteristics that distinguish them from traditional bilayer membranes. Unlike diether membranes, which consist of two separate lipid leaflets, tetraether lipids span the entire membrane, forming a monolayer structure. Pure tetraether sample exhibited notable d-spacing increases at higher humidity levels, ranging from 51 Å at 80 % relative humidity (RH) to 56 Å at 95 % RH. At high humidity, these values were comparable to D/T mixtures, whereas at lower humidity, they more closely resembled those of diether membranes **(Figure 4)**. These results are consistent with earlier findings from the Winter and Chong teams, which reported d-spacing values of 56 Å at 74 °C for *Sulfolobus acidocaldarius* polar lipid fractions using Small-Angle X-ray Scattering (SAXS) and High-Pressure FT-IR Spectroscopy (Chong et al., 2003). However, a smaller d-spacing was observed for diether membranes (ca. 50 Å). This result was unexpected, as the slip plane between their lipid layers was anticipated to result in thicker membranes compared to the tetraether membranes. Membrane thickness measurements provided further insight. Calculated head-to-head distances (d_b_) indicated a thickness of 39 Å for diether membranes and 40 Å for tetraether membranes **(Figure 8A)**. These results align with molecular dynamics (MD) simulations predicting 38 Å for diether diphytanyl phosphatidylcholine membranes and 40 Å for acyclic tetraether phosphatidylcholine membranes at 25 °C (Castro et al., 2016). SAXS intensity profiles corroborated these findings, with Gibbs-Luzzati membrane thicknesses of 38.5 Å for GDGT molecules bearing phosphatidylglucose and phosphatidylglycerol polar headgroups (Bhattacharya et al., 2024).

Remarkably, NSLD profiles revealed a small “trough” at the bilayer center (Z = 0 Å) across all membrane samples, including those rich in tetraether lipids (T and D/T) **(Figure 7)**. This feature, attributed to the density of hydrogen atoms at terminal methyl groups, is typically associated with a slip plane (Kucerka et al., 2009; Misuraca et al., 2023). However, the presence of this trough in tetraether-rich membranes was unexpected, as tetraethers span the entire membrane and lack a classical bilayer mid-plane interface. We hypothesized that this trough resulted from the shorter distance between the two methyl groups at the center of the hydrophobic chains. Due to the head-to-head condensation of two diether molecules to generate the tetraether lipid, the distance is one carbon shorter, potentially creating a localized electron density defect responsible for the observed trough (Zeng et al., 2022). Notably, this interpretation differs from electron density (ED) profiles derived from MD simulations of tetraether membranes, which do not exhibit such a feature (Kučerka et al., 2008). The discrepancy likely arises from fundamental differences in how neutrons and electrons interact with molecular structures. The slightly greater thickness of tetraether membranes may reflect the influence of their larger polar headgroups compared to diethers **(Figure 8)**. Since membrane thickness measurements (d_b_) reflect head-to-head distances, variations in thickness might primarily arise from differences in headgroup composition rather than the lipid core. Furthermore, the presence of a “trough” in NSLD profiles for both diether and tetraether membranes suggests that diether membranes might have a smaller leaflet interface than previously described **(Figure 7)**. This closer proximity of terminal methyl groups in diether membranes could enable interactions such as hydrogen bonding or Van der Waals attractions, contributing to greater stability. To further investigate these structural features, NSLD-derived density profiles of hydrocarbon chains and polar head groups were analyzed **(Figure 9)**. At 8 % D_2_O contrast, the water density profile remains approximately zero, allowing for a clear assessment of the contributions from different membrane components. The density profiles confirm that the terminal methyl groups of diether lipids are responsible for the “trough” at the membrane midplane. Interestingly, in tetraether membranes, this contribution is absent, while CH₂ groups primarily influence the NSLD profiles at Z = ± 10 Å. In contrast, for mixed D/T membranes, both CH₂ and CH₃ groups contribute to the hydrocarbon density profile, confirming the presence of both lipid types in the expected proportions. The combined analysis of NSLD and hydrocarbon chain distributions provides deeper insights into the molecular packing and stability of extremophile-derived lipids.

**Figure 9:**
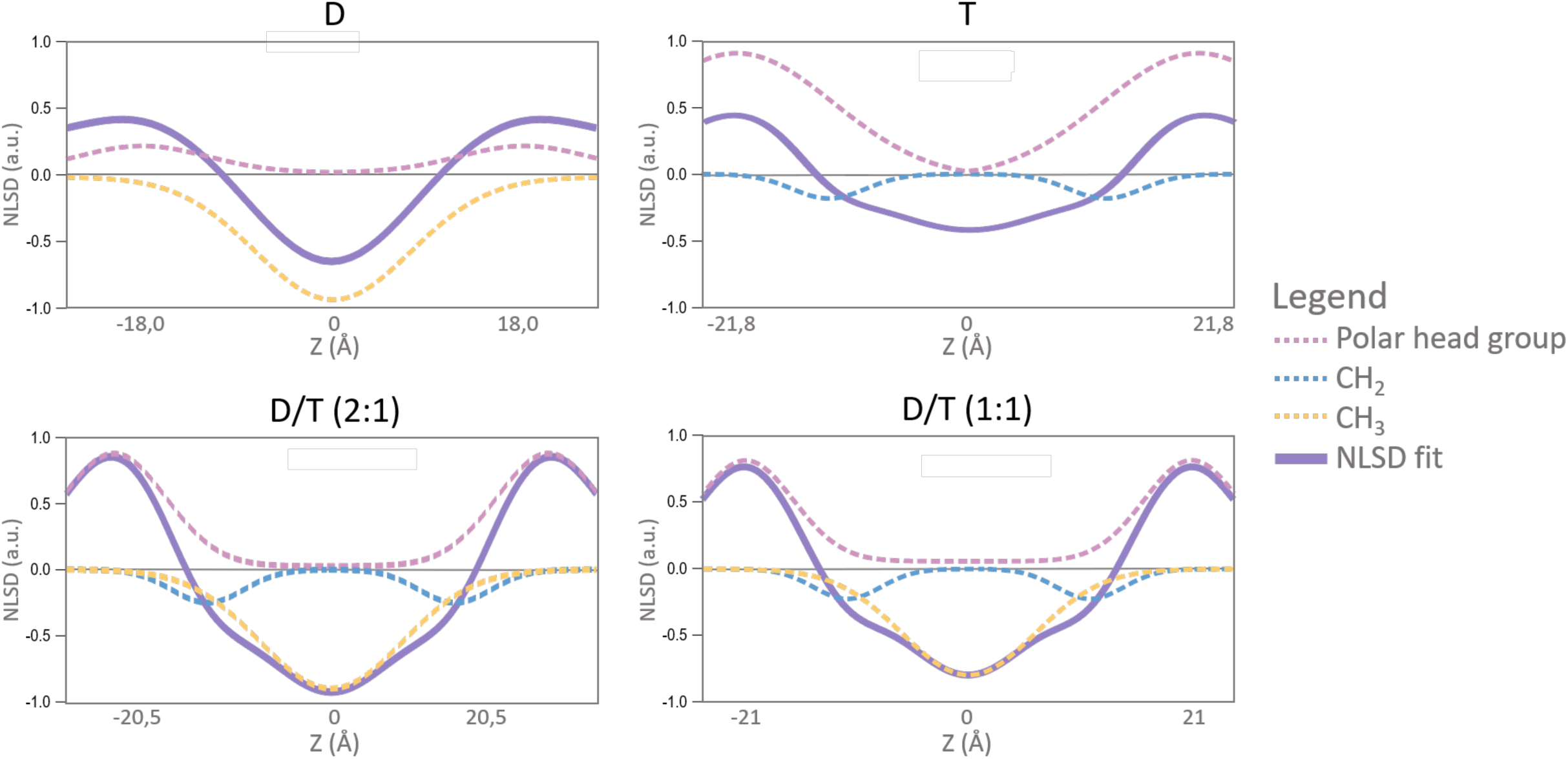
Density profiles of water, headgroups and methyl groups in 95 % hydrated membranes composed of: D= diether sample, T=tetraether sample, D/T=mixture of diether and tetraether sample at different molar ratios (1:1) or (2:1). Indicated Z position corresponds to the center of polar head group position for each sample mixture.

Cyclopentane rings in tetraether lipids also play a critical role in membrane properties. These rings, integrated into the hydrophobic core, increase membrane thickness and rigidity. MD simulations demonstrated that membrane thickness increases with ring number: ringless membranes’ thickness measured 40 Å, while membranes with four rings reached 42.6 Å (Castro et al., 2016). Similarly, calditol-GDGT membranes exhibited a decrease in thickness from 62.7 Å for ringless molecules to 45.8 Å for molecules with eight rings (Gabriel and Lee Gau Chong, 2000; Untersteller et al., 1999). Including cyclopentane rings enhances lipid packing density and thermal stability, providing a structural advantage under extreme environmental conditions. The tetraether lipid mixture used in the present study involve a mixture of lipids comprising cyclized tetraethers with two glycerols moieties and cyclized tetraethers bearing one glycerol and one calditol moiety. Furthermore, the lipid series included tetraethers with varying ring numbers, from 0 to 6. Thus, it is impossible to distinguish the relative contributions of individual ring configurations. However, it is interesting to note that the membrane thickness measured in the present study for mixed ringless and ringed tetraether lipids is closer to that measured and calculated for membranes composed of ringless tetraether lipid molecules. In summary, these findings underscore the unique structural adaptability of tetraether membranes. Tetraether lipids, with their spanning monolayer structure, cyclopentane rings, and adaptable hydrocarbon chains, demonstrate remarkable resilience under extreme conditions.

## Conclusion

Our comprehensive analysis highlights the critical role of the nature of phospholipid polar headgroups on membrane stability, especially under extreme environmental conditions. Second, our results show that lipid mixtures are essential for membrane stability and adaptability. Third, contrary to expectations, our results show that diethers may be key molecules for the structure and stability of archaeal monolayer membranes. Neutron diffraction and NSLD profiling demonstrated that tetraether membranes exhibit greater rigidity and organization in high-temperature environments compared to diether membranes. The stability of tetraether-containing membranes, as shown by consistent NSLD profiles across temperature variations **(Figure 6 and 7)**, parallels the stability seen in halophilic archaeal membranes exposed to high salt concentrations (Tenchov et al., 2006). This finding highlights the extraordinary resilience of natural lipids, especially those from hyperthermophilic Archaea, under extreme conditions.

Crucially, our study revealed that membranes composed of mixed diethers and tetraethers are more organized and stable than those formed by pure lipid systems. This structural stability, maintained across varying temperatures and humidity levels, underscores the advantage of lipid diversity in hyperthermophilic Archaea. While most hyperthermophiles possess a mixture of diethers and tetraethers, some species synthesize only one type of lipid. Our findings indicate that while these single-lipid membranes remain viable and structured at high temperatures, they lack the robustness of mixed-lipid systems. This suggests that additional membrane components may contribute to increased stability. For instance, in *Aeropyrum pernix* **(Figure 1)**, which predominantly features diether lipids with a unique C_25_ isoprenoid hydrocarbon chain, increased stability is observed compared to the typical C_20_ diether lipids (Gmajner et al., 2011; Kejžar et al., 2024). The role of apolar lipids in bilayer membrane stability has also been documented (Cario et al., 2015), with studies showing that polar polyisoprenoids aid in the adaptation of Archaea to extreme temperatures and pressures (LoRicco et al., 2020). Archaea that synthesize mostly tetraethers, also adapt to environmental stress by adjusting the degree of lipid cyclization, which influences membrane fluidity and stability under pH and thermal stress (Boyd et al., 2013; Cario et al., 2015). Alterations in hydrocarbon chains, including the formation of double bonds and cyclopentane or cyclohexane rings, help to precisely adjust membrane properties in response to environmental changes (Chong, 2010).

The ability of mixed lipid membranes to maintain integrity under extreme conditions offers valuable insights into the mechanisms of membrane stabilization and opens potential avenues for designing resilient synthetic lipid systems. Archaeal tetraether lipids have been shown to stabilize fluid diester liposomal membranes, suggesting potential applications in medical treatments, such as hyperthermia-induced drug release for cancer therapy (Ayesa and Chong, 2020). Future research incorporating X-ray scattering, molecular dynamics simulations, and computational modeling will deepen our understanding of the structural intricacies of archaeal lipid membranes, particularly the interaction between lipid composition and environmental stressors. Additionally, exploring how mixed membranes respond to other stress conditions, such as salinity and pH, is essential, given that many Archaea are multi-extremophiles.

## Acknowledgments

The authors wish to thank the ANR for financial support through grant ANR-17-CE11-0012 ArchaeoMembranes, and the Institut Laue-Langevin (Grenoble) for beamtime allocation (exp. 8-02-991 and exp. 8-02-1046) and technical support. All neutron data are publicly available on ILL servers and are accessible with the following DOIs: 10.5291/ILL-DATA.8-02-991 and 10.5291/ILL-DATA.8-02-1046. We are grateful to Thomas Hauss for sharing his Igor Pro procedure for the analysis of reciprocal space maps. MS was supported by a PhD grant from the French Ministry of Research. MT was supported by the Life Grant to SVA from the Volkswagen Foundation (Nr Az 96727).

## Author contributions

Conceptualization, P.M.O.; Methodology, M.S., M.T., P.S., S.F. and Y.L.; chemical Analysis, M.S., P.S., S.F. and Y.L; Neutron data acquisition and analysis, M.S., B.D., J.P. and P.M.O.; Original Draft Preparation, M.S. and P.M.O.; Review & Editing, M.S., P.M.O., S.F., P.S., J.P., S.V.A., and B.D.; Supervision, S.F., Y.L. and P.M.O.; Project Administration, P.M.O.; Funding Acquisition, P.M.O. and S.V.A.

## Supplementary

### Materials and Methods: Diether lipid purification

Dried cell pellets were extracted with 40 mL of a monophasic mixture of methanol (MeOH)/trichloromethane (TCM)/purified water (1:2.6:0.16; v/v/v) using a sonication probe for 10 min. After centrifugation (4000x rpm, 5 min), the supernatant was collected, and the extraction procedure was repeated twice. The supernatants were pooled and the solvents were removed under reduced pressure using a rotary evaporator. Finally, the solvent extract was solubilized using a mixture of MeOH/TCM (1:1; v/v), transferred into a 2 ml vial, and the solvents removed under an N_2_ stream. Extracted intact polar lipids (IPLs) were kept at -20 °C until lipid purification.

A two-dimensional thin-layer chromatography purification step of the IPL extract from *P. furiosus* was performed using silica gel pre-coated glass-backed plates (60 Å silica-gel, 20 cm x 20 cm plates). Polar lipids containing IPL from *P. furiosus* were separated by first developing the plate using the 1D mobile phase mixture TCM/MeOH/water (75:25:2.5; v/v/v) for 45 min in the first direction. After allowing sufficient time for drying, the plate was developed, at right angles to the first development using the 2D mobile phase TCM/MeOH/acetic acid/water (80:9:12:2; v/v/v/v) for 1.5 h (Christie, 2011). All components were detected using iodine and collected by scrapping the silica gel zone having the desired retention factor (Rf) zone. Purified lipids were extracted from the silica gel with a mixture of TCM/MeOH (1:1; v/v).

The composition of each Rf zone recovered was verified by HPLC-ESI-MS using an Inertsil Diol column (2.1 mm x 250 mm; 5 µm; GL Sciences) equipped with a pre-column with the same stationary phase. The separation was carried out in solvent gradient mode including a mobile phase solvent A (isopropanol -IPA-/water/formic acid/aqueous ammonia 88:10:0.12:0.04; v/v/v/v) and solvent B (*n*-heptane -*n*-C_7_-/IPA/formic acid/aqueous ammonia (79:20:0.12:0.04; v/v/v/v). An Agilent 1100 binary HPLC pump equipped with an autosampler and a thermostated oven set at 30 °C was used. Chemstation software (version Rev. 3.01.01.SR1) was used to control the HPLC analyses. IPL were eluted at a constant flow rate of 0.2 mL/min from 100% solvent B to 66% solvent B in 18 min, then maintained for 12 min, followed by 35% solvent B in 15 min, maintained for 15 min, and finally back to 100% solvent B in 2 min, followed by a 20 min stabilization period of the column (modified after (Peterse et al., 2011)). An Esquire 3000^Plus^ or an HCT ion trap mass spectrometer (Bruker) equipped with an electrospray ionization (ESI) source used in positive and negative modes were used. The conditions for the MS analyses were as follows: nebulizer pressure 30 psi, cone tension 40 V, drying gas (N_2_) flow 8 L/min and temperature 340 °C, capillary voltage 5 kV (negative mode) and -4 kV (positive mode), mass range *m/z* 500–2000 (Esquire 3000^Plus^) or m/z 500-3000 (HCT). Mass spectra (protonated -positive mode- and deprotonated -negative mode-) molecular ion masses of IPL were analyzed using Bruker Data Analysis software (version 4.2).

### Materials and Methods: Core lipid characterization

Aliquots of the isolated di-or tetraether polar lipids were hydrolyzed in Pyrex^TM^ tubes with PTFE caps. Polar head groups were removed using acid methanolysis (1.2 N HCl in MeOH) at 90 °C for 3 h). Once back to room temperature, the solvent, and excess reagent were removed under reduced pressure, MeOH being added several times to facilitate the removal of HCl and water. The dried crude mixture was extracted using n-C7/IPA (99:1, v/v) to recover the core lipids. In the case of the diether lipids from *P. furiosus*, the composition of the sample was controlled by HPLC-MS using the same binary pump module and ion trap mass spectrometers as described above for polar lipid analysis. The following conditions were used: HPLC column: Zorbax Sil (4.6 x 250 mm, 0.4 ml/min) connected to a pre-column with the same stationary phase, mobile phase: n-C7/IPA 95:5 (isocratic elution mode), MS source: atmospheric pressure chemical ionization source (APCI) used in the positive mode. Conditions for MS analyses were: nebulizer pressure 43.5 psi, APCI temperature 420 °C, drying temperature 350 °C, drying gas (N_2_) flow 5 L min^-1^, capillary voltage -2 kV, corona 4 mA, scan range *m/z* 500–2000 (Esquire 3000^Plus^) or *m/z* 500-3000 (HCT). The sole core lipid detected was dialkylglycerol diether (DGD).

The same analytical procedure was used for the analysis of the core lipids from *S. acidocaldarius*, except that a Zorbax RX-Sil narrow bore HPLC column (2.1 x 150 mm, 5mm; Agilent) and a mixture of n-C7/IPA (98.5:1.5 v/v) as mobile phase (isocratic elution) with a flow rate of 0.2 ml/min were used. The core lipids detected comprised glycerol dialkylglycerol tetraethers (GDGT) and calditol-GDGT having 0 to 6 cyclopentane rings.

**Figure S1:**
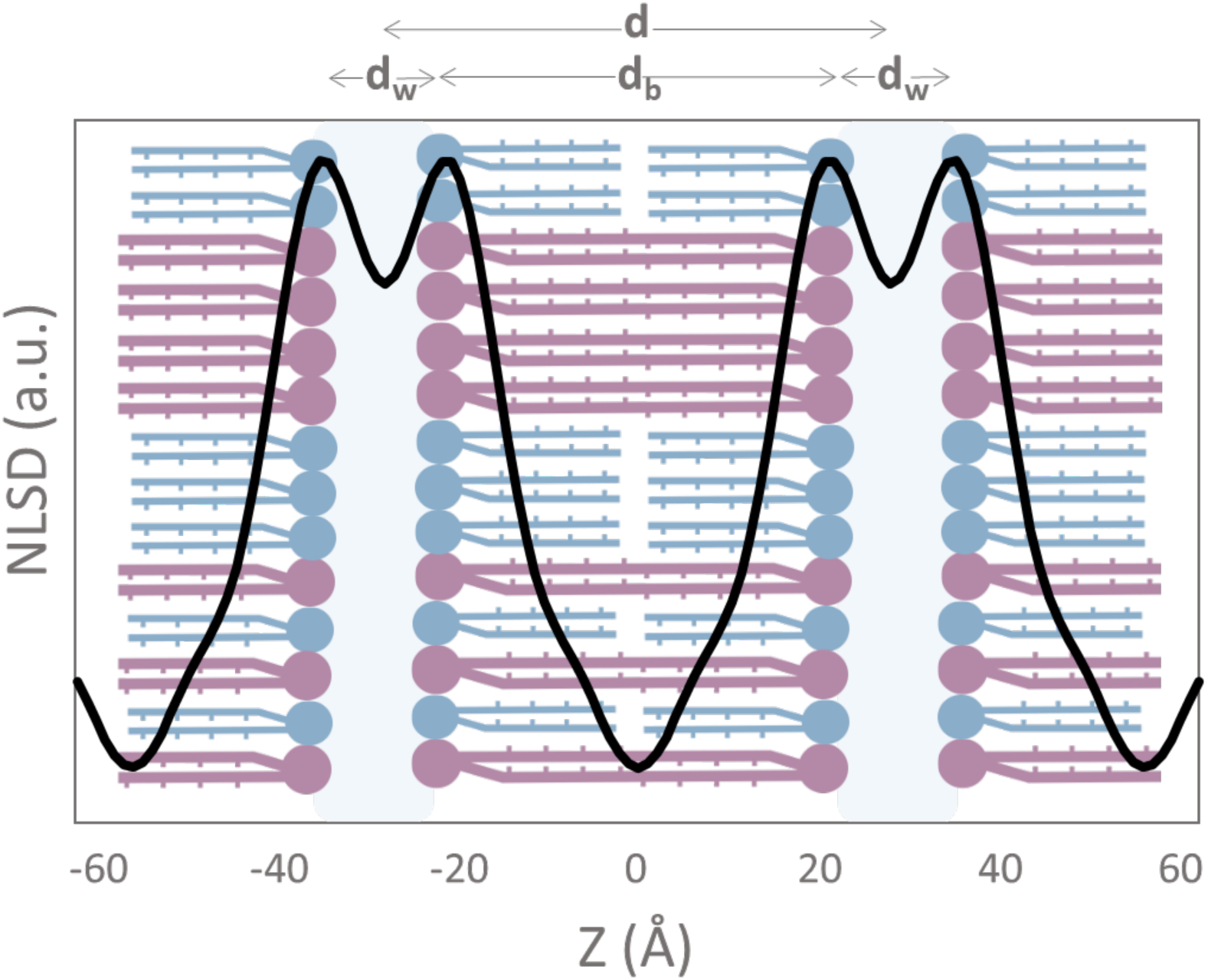
Graphic example of Neutron Scattering Length Density (NSLD) profiles. The bilayer thickness d_b_ corresponds to the center-to-center distance between headgroups as described in the methods. The water layer thickness d_w_ is calculated with d-d_b_, d being the d-spacing. D/T (1:1) 8 % D_2_O at 95 % RH, 70 °C.

**Table S1:**
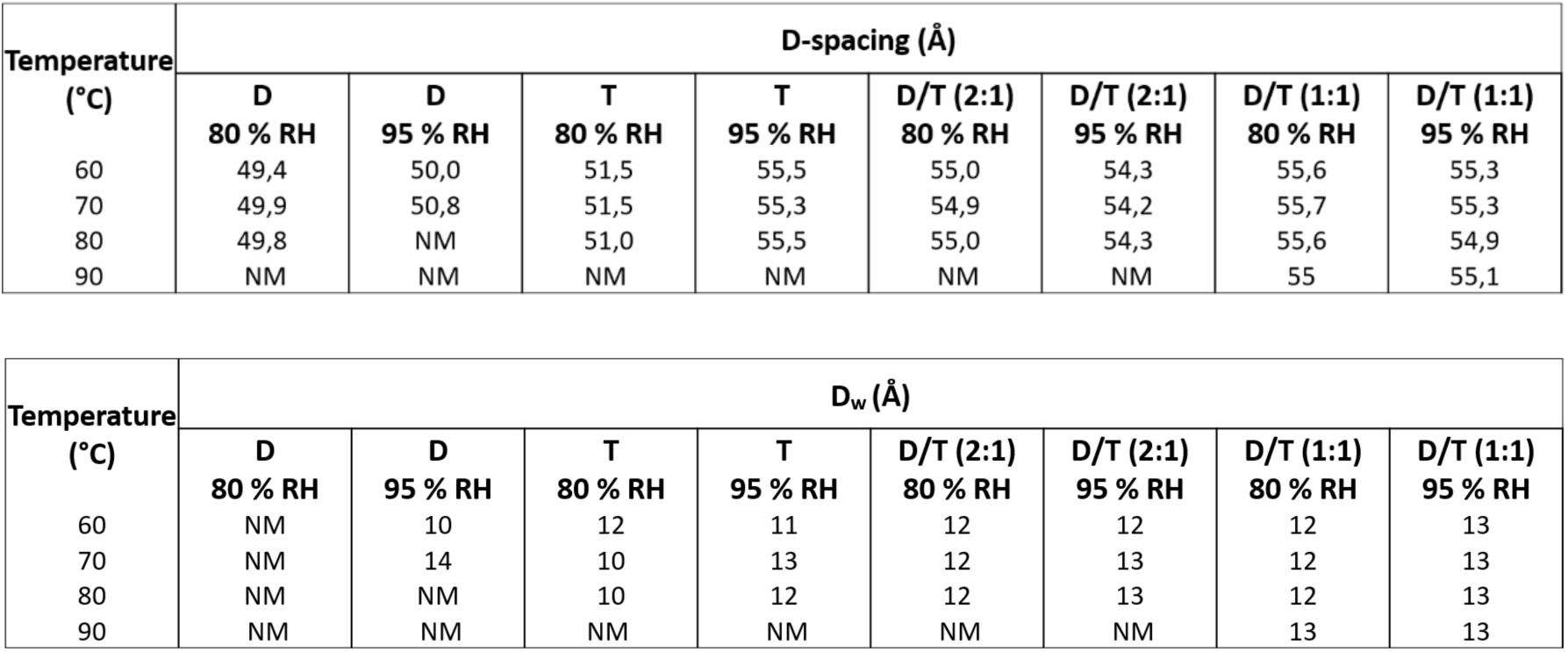
Recap table of all measurements for all tested samples. The bilayer thickness d_b_ corresponds to the center-to-center distance between headgroups as described in the methods. The water layer thickness d_w_ is calculated with d-d_b_, d being the d-spacing. D= diether sample, T=tetraether sample, D/T=mixture of diether and tetraether sample at different molar ratios (1:1) or (2:1). NM= not-measured.

## Notes

### Competing Interest Statement

The authors have declared no competing interest.

